# Replicating bacterium-vectored vaccine expressing SARS-CoV-2 Membrane and Nucleocapsid proteins protects against severe COVID-19 disease in hamsters

**DOI:** 10.1101/2020.11.17.387555

**Authors:** Qingmei Jia, Helle Bielefeldt-Ohmann, Rachel Maison, Saša Masleša-Galić, Richard Bowen, Marcus A. Horwitz

## Abstract

An inexpensive readily manufactured COVID-19 vaccine that protects against severe disease is needed to combat the pandemic. We have employed the LVS Δ*capB* vector platform, previously used successfully to generate potent vaccines against the Select Agents of tularemia, anthrax, plague, and melioidosis, to generate a COVID-19 vaccine. The LVS Δ*capB* vector, a replicating intracellular bacterium, is a highly attenuated derivative of a tularemia vaccine (LVS) previously administered to millions of people. We generated vaccines expressing SARS-CoV-2 structural proteins and evaluated them for efficacy in the golden Syrian hamster, which develops severe COVID-19 disease. Hamsters immunized intradermally or intranasally with a vaccine co-expressing the Membrane (M) and Nucleocapsid (N) proteins, then challenged 5-weeks later with a high dose of SARS-CoV-2, were protected against severe weight loss and lung pathology and had reduced viral loads in the oropharynx and lungs. Protection by the vaccine, which induces murine N-specific interferon-gamma secreting T cells, was highly correlated with pre-challenge serum anti-N TH1-biased IgG. This potent vaccine against severe COVID-19 should be safe and easily manufactured, stored, and distributed, and given the high homology between MN proteins of SARS-CoV and SARS-CoV-2, has potential as a universal vaccine against the SARS subset of pandemic causing β-coronaviruses.

---

The ongoing pandemic of COVID-19, caused by severe acute respiratory syndrome coronavirus 2 (SARS-CoV-2), has caused over 50 million cases and 1.2 million deaths as of this writing ^1^. A safe and potent vaccine that protects against severe COVID-19 disease is urgently needed to contain the pandemic. Ideally, such a vaccine would be safe, inexpensive, rapidly manufactured, and easily stored and distributed, so as to be available quickly to the entire world population.

Previously, our laboratory developed a versatile plug-and-play Single Vector Platform Vaccine against Select Agents and Emerging Pathogens wherein a single live multi-deletional attenuated *Francisella tularensis subsp. holarctica* vector, LVS Δ*capB*, is used to express recombinant immunoprotective antigens of target pathogens ^2,3^. The LVS Δ*capB* vector was derived via mutagenesis from Live Vaccine Strain (LVS), a vaccine against tularemia originally developed in the Soviet Union via serial passage and subsequently further developed and tested in humans in the USA ^4,5^. As with wild-type *F. tularensis*, LVS is ingested by host macrophages via looping phagocytosis, enters a phagosome, escapes the phagosome via a Type VI Secretion System, and multiplies in the cytoplasm ^6–8^. While much more attenuated than LVS, the LVS Δ*capB* vector retains its parent’s capacity to invade and multiply in macrophages ^9^. Using this platform technology, we have developed exceptionally safe and potent vaccines that protect against lethal respiratory challenge with the Tier 1 Select Agents of four diseases *–* tularemia, anthrax, plague, and melioidosis ^2,3^. These vaccines induce balanced humoral (antibody/neutralizing antibody in the case of anthrax toxin) and cell-mediated immune responses (polyfunctional CD4+ and CD8+ T-cells) against key immunoprotective antigens of target pathogens ^3^. We have now used this platform to develop a COVID-19 vaccine.

SARS-CoV-2 has four structural proteins – the Spike (S) glycoprotein, Membrane (M), Envelope (E), and Nucleocapsid (N) proteins. Virtually all COVID-19 vaccines in development have focused on the S protein, which mediates virus entry into host cells via the Angiotensin Converting Enzyme 2 (ACE2) receptor ^10,11^. These vaccines have been tested for efficacy most prominently in the rhesus macaque model of COVID-19. However, this is primarily a model of asymptomatic infection or mild disease, as animals typically do not develop either fever or weight loss; hence, vaccine efficacy in the rhesus macaque is quantitated primarily in terms of the vaccine’s impact on viral load rather than on clinical symptoms. In contrast, the golden Syrian hamster develops severe COVID-19 disease, akin to that of hospitalized humans ^12^, including substantial weight loss and quantifiable lung pathology.

Herein, we have employed the LVS Δ*capB* vector platform to construct six COVID-19 vaccines expressing one or more of all four structural proteins of SARS-CoV-2 (S, SΔTM, S1, S2, S2E, and MN) and tested the vaccines for efficacy, administered intradermally (ID) or intranasally (IN), against a high dose SARS-CoV-2 respiratory challenge in hamsters. We show that the vaccine expressing the MN proteins, but not the vaccines expressing the S protein or its subunits in various configurations, is highly protective against severe COVID-19 disease including weight loss and lung pathology, and that protection is highly correlated with serum anti-N antibody levels.

## Construction and verification of rLVS Δ*capB*/SCoV2 vaccine candidates

We constructed six recombinant LVS Δ*capB* vaccines (rLVS Δ*capB*/SCoV2) expressing single, subunit or fusion proteins of four SARS-CoV-2 structural proteins: S ^13^, E, M, and N **(Fig. 1A)**. The S protein is synthesized as a single-chain inactive precursor of 1,273 residues with a signal peptide (residue-15) and processed by a furin-like host proteinase into the S1 subunit that binds to host receptor ACE2 ^10^ and the S2 subunit that mediates the fusion of the viral and host cell membranes. S1 contains the host receptor binding domain (RBD) and S2 contains a transmembrane domain (TM) **(Fig. 1B, top panel).** We constructed rLVS Δ*capB*/SCoV2 expressing S (stabilized) and, so as to express lower molecular weight constructs, SΔTM, S1, S2, and the fusion protein of S2 and E (S2E), and additionally, a vaccine expressing the fusion protein of M and N (MN) (**Fig. 1B,** bottom panels). A 3FLAG-tag was placed at the N-terminus of the S, SΔTM, S1, and MN proteins. The antigen expression cassette of the SARS-CoV-2 proteins was placed downstream of a strong *F. tularensis* promoter (Pbfr) and a Shine-Dalgarno sequence (**Fig. 1B**) that we have used successfully to generate potent vaccines against *F. tularensis, Bacillus anthracis*, *Yersinia pestis*, and *Burkholderia pseudomallei*.

**Fig. 1.**
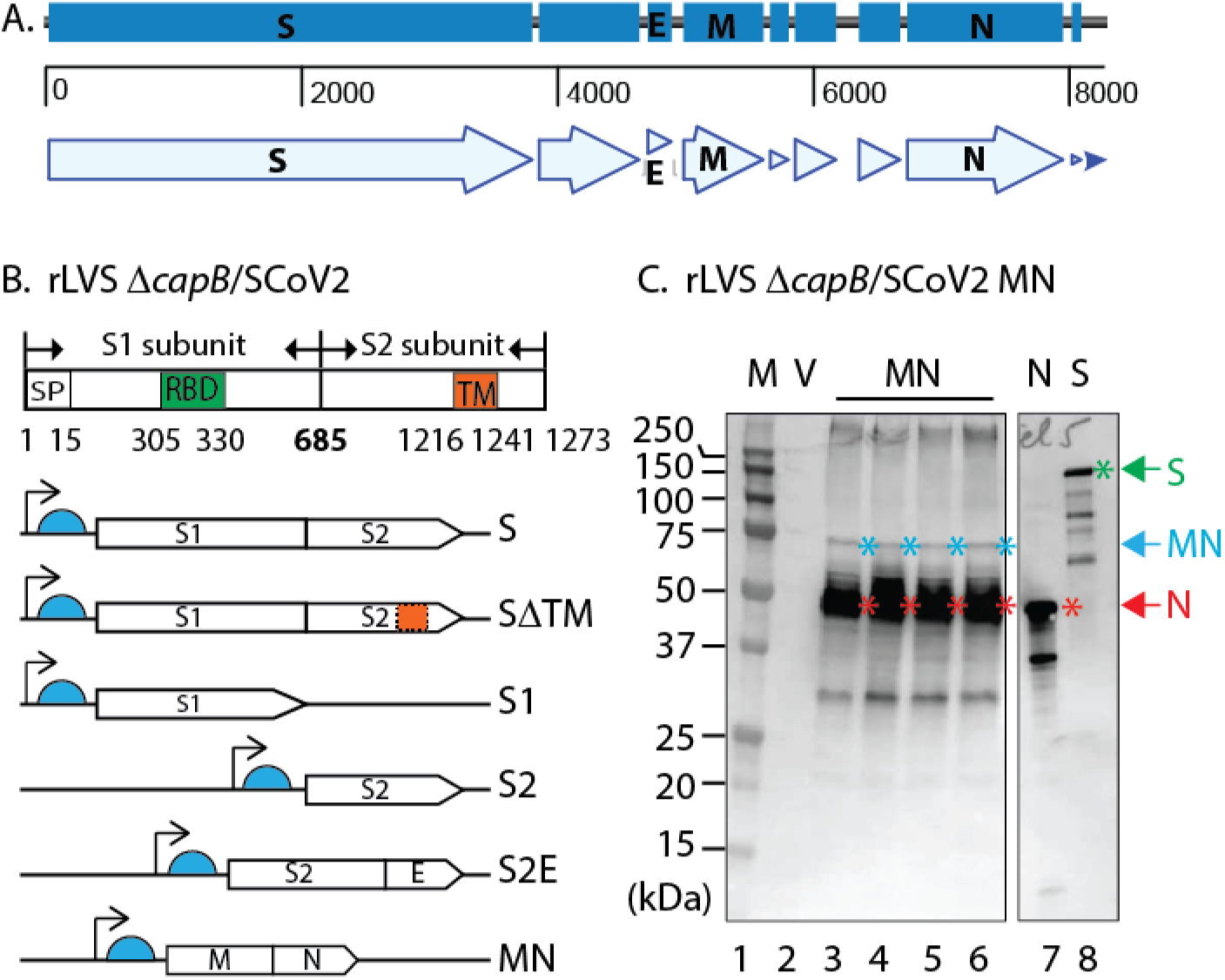
Construction of rLVS Δ*capB*/SARS-CoV-2 vaccines. **A.** Schematic of SARS-CoV-2 genomic region encoding four major structural proteins, Spike (S) glycoprotein, Envelope (E), Membrane (M), and Nucelocapsid (N) protein. **B**. Diagram of S protein and the antigen expression cassettes for S, SΔTM, S1, S2, fusion protein of S2 and E (S2E) and fusion protein of M and N (MN) downstream of the *F. tularensis bacterioferritin* (FTT1441) promoter (Pbfr) (thin black arrow) and Shine-Dalgarno sequence (light blue half circle). SP, signal peptide for S protein; RBD, receptor binding domain; and TM, Transmembrane domain. **C.** Protein expression of rLVS Δ*capB*/SCoV2 MN. Total bacterial lysates of 4 clones of rLVS Δ*capB*/SCoV2 MN (lanes 3-6, as indicated at the bottom of the lower panel) were analyzed by SDS-PAGE and Western blotting with an anti-SARS-CoV-1 guinea pig polyclonal antibody (BEI Resources, NR-10361), which readily detected the full length MN (~ 75 kDa, less abundant), indicated by blue asterisks to the right of the protein bands, and the highly abundant breakdown product N (~ 46 kDa) protein, indicated by red asterisks to the right of the protein bands. The anti-SARS-CoV-1 guinea pig polyclonal antibody also detected the N (red arrow and asterisk) and S (green arrow and asterisk) proteins of SARS-CoV-1 (lanes 7 and 8), which served as positive controls. V, LVS Δ*capB* vector (lane 2). The sizes of the molecular weight markers (M) are labeled to the left of the panels.

All six rLVS Δ*capB*/SCoV2 vaccine candidates, abbreviated as S, SΔTM, S1, S2, S2E, and MN, expressed the recombinant proteins from bacterial lysates. As shown in **Fig. 1C**, three protein bands – a minor 75 kDa, a major 46 kDa, and a minor 30 kDa band – were detected from lysates of 4 individual clones of the MN vaccine candidate (**Fig. 1C, lanes 3-6**), but not from the lysate of the vaccine vector (lane 2) by Western blotting using guinea pig polyclonal antibody to SARS-CoV, which also detected the N and S protein of SARS-CoV (lanes 7 and 8, respectively). The 75-, 46-, and 30-kDa protein bands represent the full-length MN, the N, and degradations of the MN protein. The S, SΔTM, S1, S2, and S2E proteins were also expressed by the rLVS Δ*capB*/SCoV2 vaccines, as evidenced by Western blotting analysis using monoclonal antibody to FLAG to detect S, SΔTM, and S1 (each with an N-terminus FLAG tag) and polyclonal antibody to SARS-CoV to detect non-tagged S2 protein **(Fig. S1, A-D).** Of note, SΔTM and S1 (**Fig. S1B**) were expressed more abundantly than the full-length S protein (**Fig. S1A**), possibly as a result of the removal of the TM domain and reduced size of the protein.

## Study of vaccine efficacy against SARS-CoV-2 challenge in the hamster model

We immunized Syrian hamsters (8/group, ½ female, ½ male) ID or IN twice, 3 weeks apart, with six rLVS Δ*capB*/SCoV2 vaccine candidates – S, SΔTM, S1, S2, S2E, and MN – singly and in combination (MN + SΔTM; MN + S1). Five weeks later, we challenged the animals with 10^5^ plaque forming units (pfu) of SARS-CoV-2 (2019-nCoV/USA-WA1/2020 strain) administered IN, and then closely monitored them for clinical signs of infection including weight loss. Animals immunized with PBS (Sham) or with the vector LVS Δ*capB* served as controls. At 1, 2, and 3 days post-challenge, oropharyngeal swabs were collected daily and assayed for viral load by plaque assay. At 3 and 7 days post-challenge, half of the animals in each group (4 animals, ½ male, ½ female) were euthanized and evaluated for lung viral load and lung histopathological changes, respectively **(Fig. 2A)**.

**Fig. 2.**
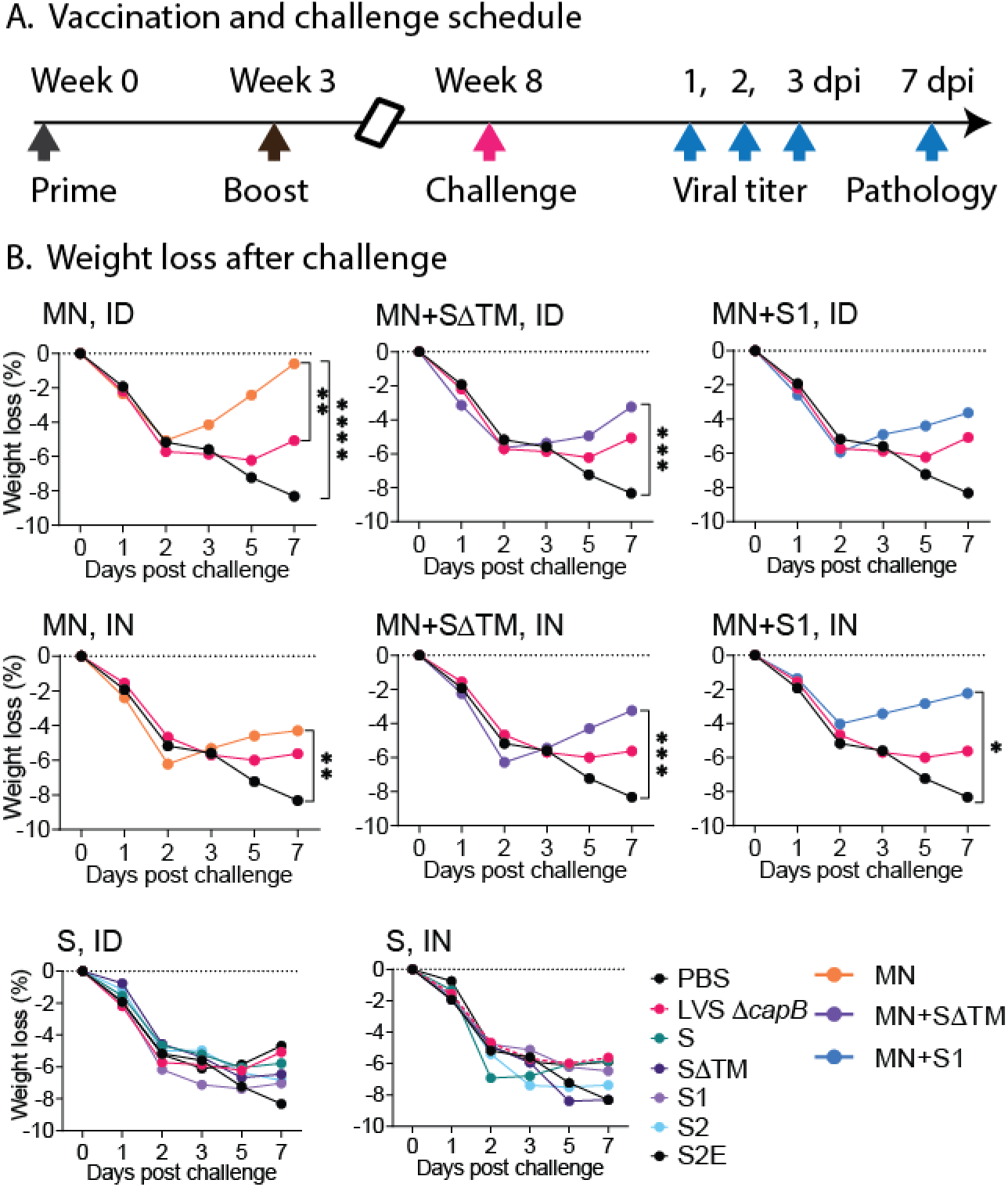
Experimental schedule and weight loss after challenge. **A.** Experiment schedule. Golden Syrian hamsters (8/group, ½ Female, ½ Male) were immunized ID or IN twice (Week 0 and 3) with rLVS Δ*capB/*SCoV2 vaccines, singly and in combination (MN+SΔTM; MN+S1); challenged IN 5 weeks later (Week 8) with 10^5^ pfu of SARS-CoV-2 (2019-nCoV/USA-WA1/2020 strain), and monitored closely for clinical signs of infection including weight loss. Single vaccines expressed the S, SΔTM, S1, S2, S2E or MN proteins, as indicated. Control animals were sham-immunized (PBS) or immunized with the vector (LVS Δ*capB*) only. All hamsters were assayed for oropharyngeal viral load at 1, 2, and 3 days post infection (dpi). Half of the hamsters (n=4/group) were euthanized at 3 dpi for lung viral load analysis and half (n=4/group) were monitored for weight loss for 7 days and euthanized at 7 dpi for lung histopathology evaluation. **B.** Weight loss post infection. Data are mean % weight loss from 1 dpi. *P<0.05; P ≤0.01; ***, P<0.001; ****, P≤0.0001 comparing means on Day 7 post-challenge by repeated measure (mixed) analysis of variance model. Sham vs. M-N: P<0.0001, ID route; P<0.01, IN route.

## MN vaccine protects against SARS-CoV-2 induced weight loss in the hamster model

As shown in **Fig. 2B**(top and middle panels), hamsters immunized either ID (top panels) or IN (middle panels) with the MN vaccine, alone or in combination with the SΔTM or S1 vaccines, were significantly protected against severe weight loss after high dose SARS-CoV-2 IN challenge [P<0.0001, P<0.01, and P<0.0001 for Sham vs MN administered ID, IN, or ID/IN (either ID or IN), respectively (Day 7) and P<0.0001 for Sham vs. all MN vaccine groups administered ID/IN] (**Table S1A**). All animals lost weight during the first 2 days after challenge; however, hamsters immunized with the MN vaccine, alone or in combination with the SΔTM or S1 vaccine, began to recover from the weight loss starting on Day 3, whereas sham-immunized animals continued to lose weight until euthanized on Day 7, by which time they had lost a mean of 8% of their total body weight. Hamsters immunized with the vector control continued to lose weight until Day 5 and then exhibited a small partial recovery, possibly reflecting a small beneficial non-specific immunologic effect as has been hypothesized for BCG and other vaccines. In contrast to hamsters immunized with the MN vaccine, hamsters immunized with the S, SΔTM, S1, S2, or S2E vaccines, administered ID or IN, were not protected against severe weight loss **(Fig. 2B, bottom panels)**.

## MN vaccine protects against severe lung pathology in the hamster model

To evaluate vaccine efficacy against SARS-CoV-2-induced lung disease, we assessed cranial and caudal lung histopathology on Day 7 post-challenge, which peaks in unvaccinated animals at this time point. As shown in **Fig. 3A and Table S2**, hamsters immunized either ID or IN with the MN vaccine, alone (MN) or in combination with SΔTM or S1, were consistently protected against severe lung pathology after high dose SARS-CoV-2 IN challenge (P< 0.0001 vs. sham-immunized hamsters for all MN containing groups, whether administered ID or IN; P<0.0001 vs. vector control for all MN groups when administered ID and P<0.01 - P<0.0001 vs. vector control for all MN groups when administered IN) (**Table S1B**). Compared with sham-immunized hamsters, the histopathology score in the cranial and caudal lungs of hamsters vaccinated with the MN vaccine was reduced on average by 71% when administered ID and 63% when administered IN. In contrast, hamsters immunized with one of the five S protein vaccines were not significantly protected against severe lung pathology whether the vaccines were administered ID or IN **(Fig. 3B)**.

**Fig. 3.**
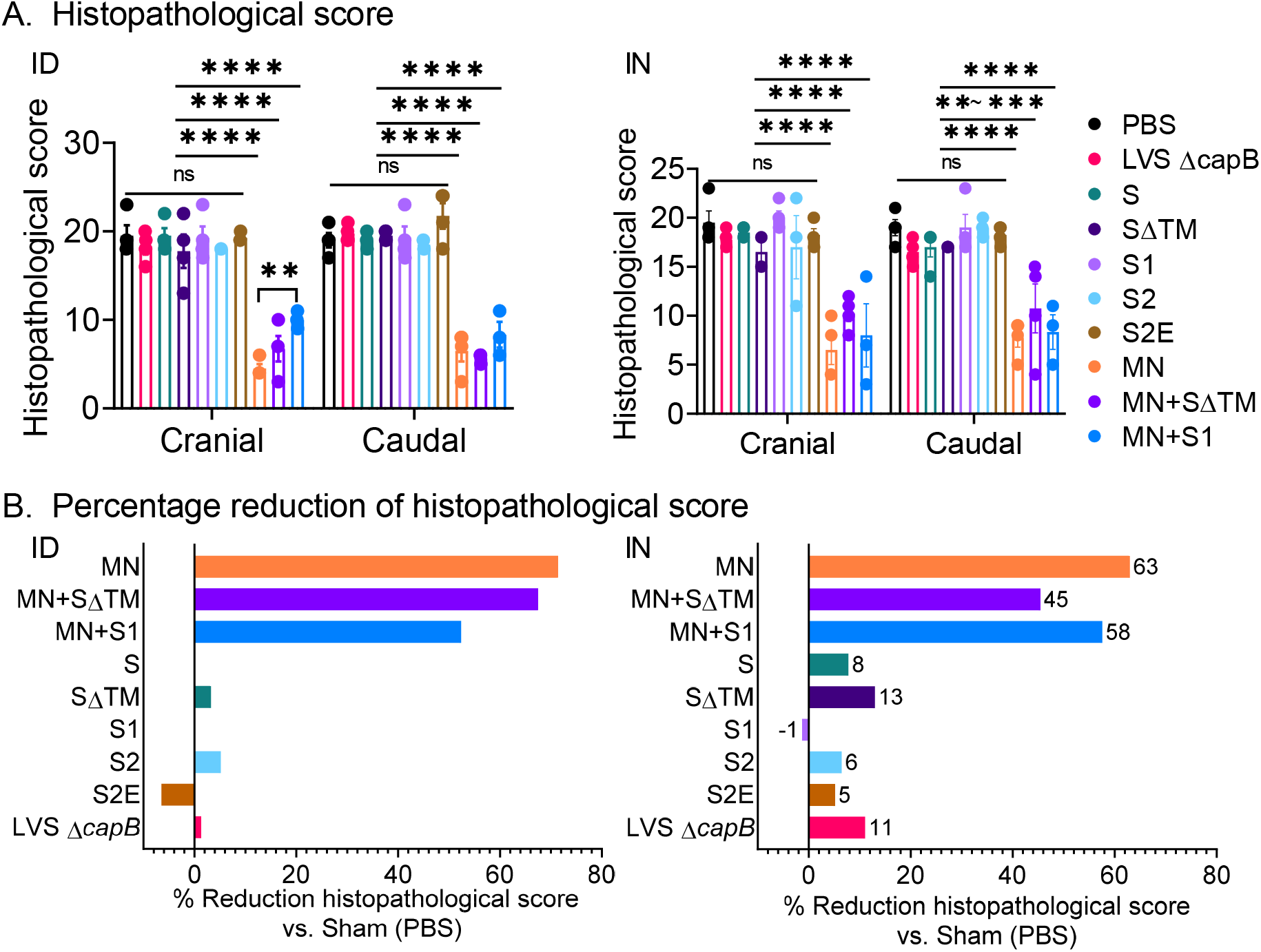
Lung histopathology on Day 7 after SARS-CoV2 IN challenge. Hamsters (n=4, ½ M, ½ F) were immunized ID or IN as described in Fig. 2 and euthanized at 7 dpi for histopathologic examination of their lungs. **A.** Cranial and caudal lung histopathology post-challenge in hamsters immunized ID (left) or IN (right) were separately scored on a 0-5 scale for overall lesion extent, bronchitis, alveolitis, pneumocyte hyperplasia, vasculitis, and interstitial inflammation; the sum of the scores for each lung are shown (mean ±SE). The histopathological score evaluation was performed by a single pathologist blinded to the identity of the groups. **P<0.01; ***, P < 0.001; ****P<0.0001 by Two-way ANOVA with Tukey’s multiple comparisons (GraphPad Prism 8.4.3); ns, not significant. **B.** The percentage reduction in the combined cranial and caudal lung histopathology score compared with Sham (PBS)-immunized animals was calculated for each vaccine.

## MN vaccine protects against SARS-CoV-2 viral replication in the oropharynx and lungs of hamsters

To examine the impact of vaccines on viral replication, we collected oropharyngeal swabs of all hamsters (n = 8/group) on Days 1, 2, and 3 post-challenge and assayed viral load by plaque assay. Hamsters immunized ID or IN with MN alone, or in combination with SΔTM or S1, showed significantly reduced viral titers in the oropharynx. Specifically, compared with sham-immunized animals, hamsters immunized ID with MN showed a 0.8±0.4, 1.0±0.4, and 1.2+0.4 log reduction (Mean ± SE) in viral load at Days 1, 2, and 3 post-challenge, respectively (P = 0.04, 0.02, and 0.004, resp.); hamsters immunized ID with MN+SΔTM or MN+S1 also showed significant reductions in viral titer compared with sham-immunized animals on Day 1 (P<0.05 for both vaccines) and, for MN+S1, on Day 3 (P<0.01) post-challenge (**Fig. 4A, left graph)**. Animals immunized IN with MN vaccines (MN, MN+SΔTM, MN+S1) also showed reduced viral load compared with sham- and vector-immunized animals on Days 1-3 post-challenge, on average 0.8±0.3, 0.8±0.3, and 0.6±0.3 logs fewer than Sham on Days 1-3 post-challenge (P <0.02 for all MN vaccines vs. Sham on Days 1 and 2 post-challenge) **(Fig. 4A, right graph).** All MN vaccines combined, whether administered ID or IN, showed mean reductions compared with Sham of 0.9±0.3, 0.6±0.3 and 0.8±0.3 logs on Days 1, 2, and 3, respectively (P< 0.01, P< 0.05, and P< 0.01, resp.). In contrast, hamsters immunized with the S protein vaccines (S, SΔTM, S1, S2, and S2E) did not show significantly reduced viral titers compared with sham-immunized animals whether the vaccines were administered ID or IN **(data not shown)**.

**Fig. 4.**
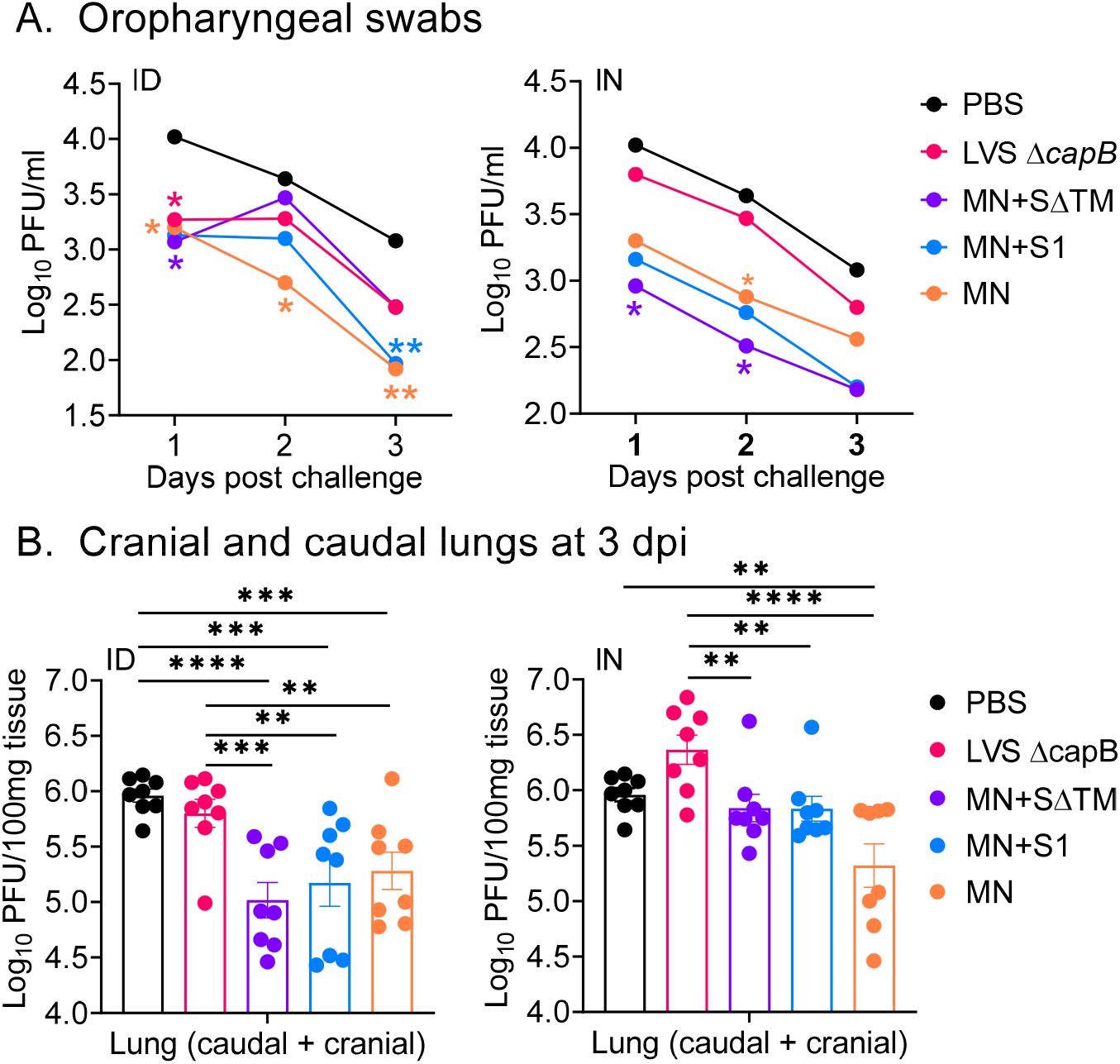
Viral load in oropharyngeal swabs and cranial and caudal lungs 3 days post-challenge. Hamsters were immunized ID or IN as described in Fig. 2. **A.** Oropharyngeal swabs were collected at 1, 2, 3 dpi and assayed for viral load by plaque assay. Data are Mean Log10 PFU per ml. *, P<0.05; **, P<0.01 (color coded to each vaccine candidate) vs. Sham (PBS) by repeated measure analysis of variance. **B.** Cranial and caudal lung homogenates were prepared at 3 dpi and assayed for viral titer. Data are mean ± SE of PFU per 100 mg of homogenized tissue, as indicated. **, P<0.01; ***, P<0.001; ****, P<0.0001 by Two-way ANOVA with Dunnett multiple comparisons (GraphPad Prism 8.4.3).

To evaluate viral replication in the lungs, we assayed cranial and caudal lungs for viral load on Day 3 post-challenge, which peaks at this time point in unvaccinated animals. Hamsters immunized ID with the MN vaccine, alone or in combination with the SΔTM or S1, showed significantly reduced viral loads in their cranial and caudal lungs compared with sham- or vector-immunized animals **(Fig. 4B, left panel)**. Hamsters immunized ID with the MN vaccines as a group showed a mean reduction of 0.8±0.1 log compared with Sham (P< 0.0001). In contrast, hamsters immunized ID with the S (S, SΔTM, S1, S2, S2E) protein vaccines did not show reduced viral loads in their cranial and caudal lungs (data not shown). Similar results were observed in hamsters immunized IN (**Fig. 4B**, right panel).

## MN expressing vaccines induce antibody to N protein with a TH1 bias

To assess antibody responses to SARS-CoV-2 proteins expressed by the vaccine, we analyzed antibodies to the RBD of the S protein and to the N protein **(Fig. 5)**. As expected, sera from sham- and vector-immunized hamsters lacked antibody to either antigen **(Fig. 5A-C)**. In contrast, sera from hamsters immunized once with the MN vaccine, alone or in combination with the SΔTM or S1 vaccine, showed high levels of N specific IgG, whether immunized ID or IN, at 3 weeks post-immunization (**Fig. 5A**), which somewhat increased at Week 8, 5 weeks after the second immunization at Week 3 (**Fig. 5B**), displaying a TH1 type bias, with IgG2 dominating the response (**Fig. 5C**). Differences in serum anti-N IgG titers between hamsters immunized with the MN vaccine, alone or in combination with S protein vaccines, and sham- or vector-immunized hamsters were highly significant at both Week 3 and Week 8 (P<0.0001) (**Fig. 5D**). Surprisingly, hamsters immunized with S protein vaccines did not show anti-RBD antibody at Week 3 (Fig. 5A), nor SARS-CoV-2 neutralizing antibody at Week 8 (data not shown). In mice immunized at Weeks 0 and 3 with second generation vaccines expressing MN in combination with S1 or SΔTM, serum obtained at Week 4 showed anti-RBD antibody as well as anti-N antibody **(Fig. S2)**. Anti-N IgG antibody displayed a TH1 type bias both in hamsters (**Fig. 5C**), where IgG2 dominated the IgG response, and in mice, where IgG2a dominated the IgG response **(Fig. S2).** This TH1 bias was also reflected by murine splenocyte secretion of IFN-γ in response to S and N peptides **(Fig. S3)**.

**Fig 5.**
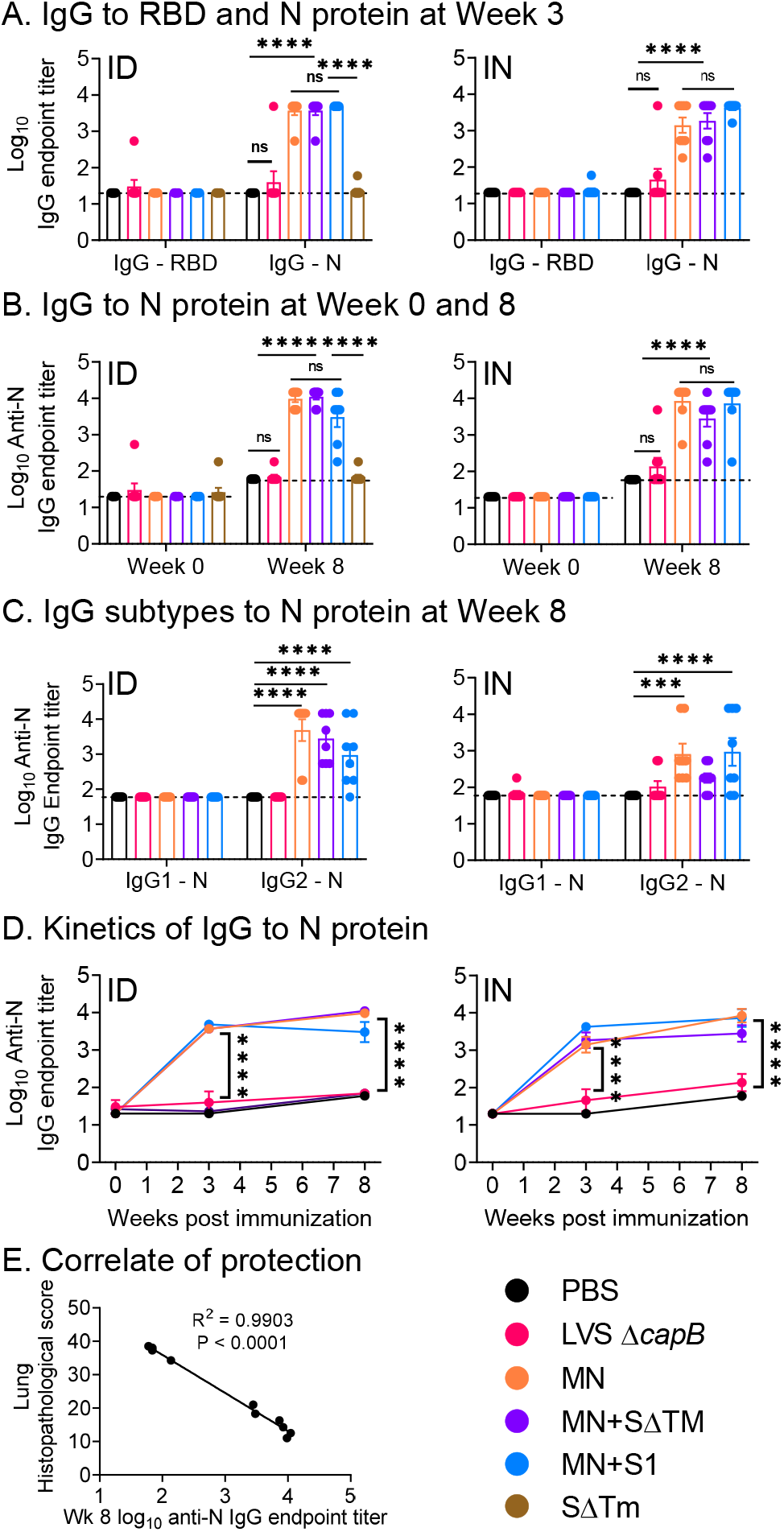
Humoral immune response and correlate of protection. Hamsters were immunized ID or IN as described in Fig. 2. **A.** Sera were evaluated for IgG specific to S RBD and N protein three weeks after a single immunization ID (left) or IN (right). **B.** Sera were evaluated for IgG specific to N protein just prior to immunization at Week 0 and just prior to challenge at Week 8 in hamsters immunized ID (left) or IN (right). **C.** Sera were evaluated for IgG subtypes (IgG1 and IgG2) specific to N protein at Week 8 in hamsters immunized ID (left) or IN (right). **D.** The antibody titers displayed in **A** and **B** are plotted over time. ***, P<0.001; ****, P<0.0001; ns, not significant by Two-way ANOVA with Tukey’s multiple comparisons (GraphPad Prism 8.4.3). **E.** Correlation between IgG N titers at Week 8 and lung histopathology score on Day 7 post-challenge (sum of cranial and caudal lung as shown in Fig. 3A).

## Serum anti-N antibody correlates with protection in hamsters

We assessed the correlation coefficient between serum anti-N IgG antibody just before challenge at Week 8 and lung (cranial + caudal) histopathological scores at Day 7 post-challenge by linear regression analysis. Anti-N antibody was highly and inversely correlated with histopathology score (R^2^= 0.9903, P< 0.0001) (**Fig. 5E**). This antibody, which does not neutralize SARS-CoV-2 (data not shown), likely is not itself protective but instead correlates with a protective T cell response such as that shown in Fig. S3.

## Discussion

We show that a replicating LVS Δ*capB*-vectored COVID-19 vaccine, rLVS Δ*capB*/SCoV2 MN, that expresses the SARS-CoV-2 M and N proteins, protects against COVID-19 disease in the demanding golden Syrian hamster model. The vaccine significantly protects against weight loss and severe lung pathology, the two major clinical endpoints measured, and significantly reduces viral titers in the oropharynx and lungs. The vaccine was protective after either ID or IN administration.

Surprisingly, of the six vaccines expressing one or more of the four SARS-CoV-2 structural proteins, only the vaccine expressing the MN proteins was protective. Such a vaccine has the potential to provide cross-protective immunity against the SARS subgroup of β-coronaviruses including potential future pandemic strains. While the S protein shows only 76% sequence identity between SARS-CoV and SARS-CoV-2, the M and N proteins each show 90% identity^14^. In an analysis of T-cell epitopes in humans recovered from COVID, the M and N antigens together accounted for 33% of the total CD4+ T cell response (21% and 11% for M and N, respectively) and 34% of the total CD8+ T cell response (12% and 22% for M and N, respectively), an amount exceeding the 27% and 26% CD4 and CD8 T cell responses, respectively, of the S protein ^15^. Hence, the MN vaccine has potential for universal protection against this group of especially severe pandemic strains.

We evaluated our vaccines in the hamster model of SARS-CoV-2 infection because of its high similarity to serious human COVID-19 disease, which likely reflects at least in part the high genetic similarity of the hamster and human ACE2 receptor – S protein interface. A modelling of binding affinities showed that the hamster ACE2 has the highest binding affinity to SARS-CoV-2 S of all species studied with the exception of the human and rhesus macaque.

In our previous studies of vaccines utilizing the LVS Δ*capB* vector platform, three immunization doses consistently yielded superior efficacy to two doses. Here, given the urgency for a COVID-19 vaccine and the desire to simplify the logistics of vaccine administration, we opted to test only two immunizations, while still maintaining a reasonably long immunization-challenge interval (5 weeks after the second immunization). Future studies will examine if three doses are superior to two and the longevity of immunoprotection.

Generally speaking, vaccine efficacy in a relevant animal model of disease is the best predictor of vaccine efficacy in humans. That non-human primates (NHPs) challenged with SARS-CoV-2 develop only mild disease or remain asymptomatic brings into question the utility of this animal model as a predictor of COVID-19 vaccine efficacy in humans. Nevertheless, most published studies on vaccine efficacy have been conducted in NHPs ^16–21^; the absence of quantifiable clinical symptoms limited these studies to measuring differences in viral load between immunized and control animals. In contrast to NHPs, SARS-CoV-2 challenged hamsters develop quantifiable clinical symptoms, especially weight loss and lung pathology, akin to humans seriously ill with COVID-19. Since the major utility of a vaccine is in preventing serious infection and death, the hamster is a highly relevant animal model for assessing COVID-19 vaccine efficacy, and studies in hamsters can be conducted at a fraction of the cost, complexity, human and facility resources, and ethical concerns of studies in NHPs.

Many types of vaccines are being developed against COVID-19 including DNA, RNA, and protein/adjuvant vaccines, non-replicating and replicating viral-vectored vaccines, whole inactivated virus vaccines, and virus-like particles. To our knowledge, ours is the only vaccine comprising a replicating bacterial vector. Replicating vaccines are among the most successful vaccines in history with a reputation for inducing comprehensive immune responses and long-lasting immunity ^22^.

Our LVS Δ*capB*-vectored vaccine platform offers several advantages including 1) low toxicity;) ability to express multiple antigens of target pathogens from two different sites (shuttle plasmid and chromosome); 3) balanced immunogenicity – B-cell and T-cell (TH1 type); 4) ease of administration by multiple routes (intradermal, subcutaneous, intramuscular, intranasal, oral, etc.); 5) no animal products in contrast to viral-vectored vaccines grown in cell culture; 6) no need for adjuvant; 7) no pre-existing immunity as with adenoviruses; 8) low cost of manufacture as extensive purification is not required, in contrast to RNA, protein/adjuvant and viral-vectored vaccines; 9) ease of large scale manufacture via bacterial fermentation in simple broth culture; and 10) after lyophilization, convenient storage and distribution at refrigerator temperatures. These last three advantages are particularly important with respect to making a COVID-19 vaccine available rapidly and cheaply to the entire world’s population.

Safety is always a major consideration in vaccine development, especially so in the case of replicating vaccines. In our vaccine’s favor, its much less attenuated parent (LVS) was already considered safe enough to justify extensive testing in humans, including recently, and it has demonstrated safety and immunogenicity ^5,23–29^. LVS has two major attenuating deletions and several minor ones ^30^. As many as 60 million Russians were reportedly vaccinated against tularemia with the original LVS strain ^31^, and over 5,000 laboratory workers in the United States have been vaccinated with the modern version of LVS by scarification ^5^. Our further attenuation of LVS by introduction of the *capB* mutation reduced its virulence in mice by the IN route by >10,000-fold ^9^. Hence, rLVS Δ*capB*/SCoV2 MN and other LVS Δ*capB*-vectored vaccines are anticipated to be exceedingly safe.

Correlates of protective immunity to COVID-19 are not well understood. Almost all of the vaccines in development are centered on generating immunity to the S protein - especially neutralizing antibody to this protein. However, neutralizing antibody alone may not be sufficient for full protection; vaccines generating strong neutralizing antibody responses against SARS-CoV were not necessarily highly protective, especially in ferrets, which exhibit SARS disease more akin to that in humans ^32,33^. T-cell responses may be as or more important. T cell responses were demonstrated to be required to protect against clinical disease in SARS-CoV challenged mice and adoptive transfer of SARS-CoV specific CD4 or CD8 T-cells into immunodeficient mice infected with SARS-CoV lead to rapid viral clearance and disease amelioration ^34^.

Our S protein vaccines were ineffective, likely due to suboptimal S protein immunogenicity, reflected by the rapid decline of antibody titer in mice and the negligible antibody neutralization titers in hamsters just before challenge (data not shown). Possibly, enhanced or alternative expression of the S protein, for example display on the bacterial surface in addition to secretion, would improve immunogenicity, as reported for the S protein of SARS-CoV ^35^. This would allow immune responses to the S protein to contribute to the already substantial protective efficacy provided by immune responses to the M and N proteins.

Our replicating bacterial vaccine expressing the M and N proteins has demonstrated safety and efficacy in an animal model of severe COVID-19 disease. If its safety and efficacy is reproduced in humans, the vaccine has potential to protect people from serious illness and death. Considering the ease with which our vaccine can be manufactured, stored, and distributed, it has the potential to play a major role in curbing the COVID-19 pandemic, thereby saving thousands of lives and more rapidly restoring the world’s battered economy.

## Acknowledgments

We thank Lauren Guilbert, Airn Hartwig, and Stephanie Porter for technical support with *in vivo* challenge experiments at CSU and Jeffrey Gornbein for assistance with statistical analyses. The following reagents were obtained through BEI Resources, NIAID, NIH: SARS-CoV-2 virus (2019-nCoV/USA-WA1/2020 strain), recombinant N, S, and RBD proteins, and guinea pig polyclonal anti-SARS coronavirus antibody.

## Author contributions

M.A.H .and Q.J. conceived the project; M.A.H. oversaw the project and obtained funding; R.B. oversaw the *in vivo* challenge experiment in hamsters; Q.J. designed molecular constructs; Q.J. and S.M.G. completed vaccine molecular construction; R.M. conducted the hamster study and assayed viral load in hamster tissues and neutralizing antibody in hamster sera. H.B-O. assessed lung histopathology. Q.J. and S.M.G assayed hamster and mouse serum antibody and mouse T cell immune responses; Q.J. processed data; Q.J. and M.A.H. wrote the manuscript. All authors reviewed the manuscript in final form.

## Data availability

All data supporting the findings of this study are available within the article and its supplementary information files or from the corresponding author upon request.

