## Supplement for "Replicating bacterium-vectored vaccine expressing SARS-CoV-2 Membrane and Nucleocapsid proteins protects against severe COVID-19 disease in hamsters"

### Materials and Methods

#### Ethics Statement

Hamsters and mice were used according to protocols approved by the Institutional Animal Care and Use Committees of Colorado State University (CSU) and UCLA, respectively.

**Cells, virus and bacteria.** Vero E6 cells (ATCC #CCL-81, Manassas, VA, USA) were grown in Dulbecco modified Eagle Medium (DMEM) with high glucose (Millipore Sigma, St. Louis, MO, USA). SARS-CoV-2 virus (2019-nCoV/USA-WA1/2020 strain) was acquired through the NIH NIAID Biodefense and Emerging Infections Research Resources Repository (BEI Resources), passaged three times in Vero E6 cells, and stocks frozen in DMEM supplemented with 10% fetal bovine serum. The virus titer was determined by plaque assay as described previously [36] and below. *F. tularensis* Live Vaccine Strain with a deletion in *capB* (LVS Δ*capB*) was constructed as described by us previously [9]. Live attenuated recombinant LVS Δ*capB* expressing SARS-CoV-2 antigens (rLVS Δ*capB*/SCoV2 and rLVS Δ*capB*::MN/SCoV2) were constructed as described below. Stocks of LVS Δ*capB* vector, rLVS Δ*capB*/SCoV2, and rLVS Δ*capB*::MN/SCoV2 vaccine candidates were prepared in broth medium. Briefly, the bacteria were inoculated in Medium T broth [37] at an initial optical density (OD) of 0.003 – 0.005 at 600 nm absorbance, grown overnight at 37°C with shaking, harvested by centrifugation at 5000x g for 20 min, washed twice with sterile normal saline, suspended in 20% glycerol – normal saline solution, and frozen in 0.25 ml aliquots at -80°C until use. The titers of bacterial stocks were determined immediately before and periodically after freezing by spotting 0.05 ml of 10-fold serial diluted bacterial suspension onto Chocolate agar plates supplemented with or without kanamycin (7.5 µg/ml); and the plates were incubated at 37°C for 3 – 5 days before colony forming units (CFU) were counted.

**Proteins, antibodies, and heat-inactivated viruses and bacteria.** We obtained the following reagents through BEI Resources: SARS coronavirus S glycoprotein with deleted TM domain, SΔTM, recombinant from Baculovirus (NR-722); SARS-CoV-2 S glycoprotein (stabilized), recombinant from Baculovirus (NR-52396; NR-52308); SARS-CoV-2 S glycoprotein RBD with C-Terminal histidine tag, recombinant from HEK293T Cells (NR-52946); SARS coronavirus N glycoprotein, recombinant from *E. coli* (NR-699); SARS-CoV-2 N protein N-terminal RNA binding domain with N-terminal histidine tag, recombinant from *E. coli* (NR-53246); SARS-CoV-2 S glycoprotein peptide array (NR-52402); SARS-CoV-2 N protein peptide array (NR-52404); heat inactivated SARS-CoV-2, isolate USA-WA1/2020 (NR-52286); and guinea pig polyclonal anti-SARS coronavirus antibody (NR-10361). Monoclonal anti-FLAG M2 HRP antibody was purchased from Millipore Sigma (St. Louis, MO). Stocks of heat-inactivated LVS Δ*capB* (HI-LVS) were prepared as described previously [9].

**Generation of rLVS Δ*capB*/SCoV2 vaccines.** Six live attenuated recombinant LVS Δ*capB* expressing SARS-CoV-2 structural proteins (rLVS Δ*capB*/SCoV2) S, SΔTM, S1 subunit, S2 subunit, or fusion proteins of S2E (S2 subunit fused to E protein) and MN (M protein fused to N) from a shuttle plasmid were constructed by electroporating a shuttle plasmid carrying a SARS-CoV-2 antigen expression cassette downstream of the *Francisella* bacterioferritin (bfr) promoter (Pbfr) into rLVS Δ*capB,* as we published previously [3]. Briefly, to construct a shuttle plasmid expressing the S protein (protein id QIH55221.1), we codon-optimized a gene (Genebank MT152824) encoding the full-length SARS-CoV-2 S protein with two stabilizing proline substitutions at the S2 fusion machinery (K986P and V987P) [38,39] for expression in LVS and had it synthesized by Atum.com. Similarly, genes encoding fusion proteins of S2E and MN, linked by a flexible linker (GGSG), were codon-optimized and synthesized by Atum.com. Subsequently, we cloned the codon-optimized DNA for S, S2E, and MN with an N-terminal 3FLAG tag into the pFNL-derived shuttle plasmid downstream of the Pbfr promoter to generate pFNL/Pbfr-N3F-S, pFNL/Pbfr-N3F-S2E, and pFNL/Pbfr-N3F-MN. We further mutated the S protein expression cassette in the pFNL/Pbfr-N3F-S plasmid and generated pFNL/Pbfr-N3F-SΔTM, pFNL/Pbfr-N3F-S1, and pFNL/Pbfr-S2 by using the QuikChange Lightning Site-Directed Mutagenesis Kit (Agilent Technologies, https://www.agilent.com) and the following primer pairs: TGAGGTTAAGGATCCACTAGCTCGTTTCAAA and TTGTTCGTATTTTCCAAGTTCTTGTAGATCTATTAAA for generating pFNL/Pbfr-N3F-SΔTM; TGAGGTTAAGGATCCACTAGCTCGTTTCAAA and TCTTGCGCGACGAGGACTATTTGTCTGTGT for generating pFNL/Pbfr-N3F-S1; and TCAGTAGCATCACAATCGATTATAGCTTATACAA and CATTACGTACCTCCTATTGTTACCTCCATTATTTA for generating pFNL/Pbfr-S2. After verifying the nucleotide sequences for the SARS-CoV-2 protein expression cassette by restriction analysis and nucleotide sequencing, we electroporated the pFNL plasmid that carries a kanamycin-resistance gene into LVS Δ*capB* electro-competent cells; selected recombinant clones (rLVS Δ*capB*/SCoV2 S, SΔTM, S1, S2, S2E, and MN) on Chocolate agar plates supplemented with kanamycin (7.5 µg/ml); and verified kanamycin-resistant clones by nucleotide sequencing of the antigen expression cassette and by Western blotting for protein expression.

**Efficacy study in hamsters.** Golden Syrian hamsters (*Mesocricetus auratus*), age 9 weeks, were purchased from Charles River Laboratories). Animals (8/group, 4 male, 4 female) were immunized i.n. or i.d twice, 3 weeks apart (Week 0 and 3), with 4 x $10^6$ CFU ID or 2 x $10^6$ CFU IN of rLVS Δ*capB*/SCoV2 S, SΔTM, S1, S2, S2E, or MN vaccine candidates. The MN vaccine was also administered in combination with the SΔTM or S1 vaccine ID (4 x $10^6$ CFU each) or IN

(1 x $10^6$ CFU each). Hamsters vaccinated with normal saline (sham) or LVS Δ*capB* (vector) served as controls. Blood was collected 1-3 days prior to each immunization and challenge to assess antibody responses; the blood was heat inactivated at 56°C for 30 min. All the animals were challenged i.n. at Week 8 with approximately $10^5$ pfu of SARS-CoV-2 (2019-nCoV/USA-WA1/2020 strain) under light anesthetized with ketamine-xylazine. Virus diluted in phosphate buffered saline (PBS) was administered via pipette into the nares (100 ul total, ~50 ul/nare); animals were observed until fully recovered from anesthesia. Virus back-titration was performed on E6 cells immediately following inoculation, confirming that hamsters received 1.8 (males) or 2.2 (females) x $10^5$ pfu. Hamsters were moved to ABSL3 6-8 days prior to challenge. Animals (n=8/group) were monitored daily post-challenge for clinical signs of infection (fever, weight loss, nasal discharge, etc.); weighed on Days 1, 2, 3, 5, and 7 post-challenge; and the oropharynx swabbed for virus titers on Days 1, 2, and 3 post-challenge. Half the animals (4 hamsters) in each group were euthanized at Day 3 post-challenge (acute phase) to assess lung (cranial and caudal lobes) virus titer, which peaks at Day 3, and the other half of each group euthanized at Day 7 (subacute phase) post-challenge to evaluate histopathology, which peaks at that time.

**Histopathology assessment.** Tissues from hamsters were fixed in 10% buffered formalin for 7 to 14 days, embedded in paraffin, and cut slides stained with hematoxylin and eosin. Slides were read by a single veterinary pathologist blinded to the identity of the vaccine groups. Cranial and caudal lung histopathology were separately scored for: overall lesion extent, bronchitis, alveolitis, pneumocyte hyperplasia, vasculitis, and interstitial inflammation, each on a 0-5 scale, and the scores for each lung summed. In addition, animals were evaluated for evidence of antibody-dependent enhancement (ADE) and eosinophilic inflammation.

**Virus assay.** Virus titration was performed on oropharyngeal swabs obtained at 1, 2, and 3 dpi and on tissue samples of cranial and caudal lungs obtained at 3 dpi by double-overlay plaque assay on Vero E6 cells as previously described [36]. Briefly, fluids or tissue homogenates were serially diluted in BA1 medium and inoculated onto confluent monolayers of Vero E6 cells seeded in 6-well cell culture plates; incubated at 37$^{o}$C for 45 min; and each well was overlaid with 2 ml of MEM containing 2% fetal bovine serum and 0.5% agarose. After 24 to 30 hours incubation at 37$^{o}$C in 5% $CO_2$, a second overlay identical to the first but containing neutral red dye was added and the infected cells continued to culture. At 48-72 hours post-infection, plaques were counted with the aid of a lightbox.

Plaque reduction neutralization assay (PRNT) was performed on heat-inactivated hamster sera as described previously [36].

**Enzyme-linked immunosorbent assay (ELISA) for anti-SARS-CoV-2 RBD and N antibody in hamster sera.** Hamster sera were assayed for IgG and subtype antibodies specific to SARS-CoV-2 S RBD and N protein antigens as described by us previously [9]. Briefly, RBD or N proteins (1 µg/ml) were diluted in carbonate/bicarbonate buffer (50 mM $NaHCO_3$, 50 mM $Na_2CO_3$) and 0.1 ml was used to coat 96-well high-binding capacity plates (Corning, NY) overnight at 4$^{o}$C. Excess antigen was removed and residual antigen blocked in Blocker Casein in PBS [Thermo Scientific] for 1 hour at room temperature. Sera at a starting dilution of 1:60 were diluted further through a three-fold series with PBS containing 1% bovine serum albumin. The diluted sera were incubated with antigens coated on 96-well plates for 90 min, then incubated for

90 min with horseradish peroxidase (HRP)-conjugated mouse anti-hamster IgG (ThermoFisher), IgG1, or IgG2 (Southern Biotech) at a dilution of 1:1000 at ambient temperature. The plates were washed three times with PBS containing 0.05% Tween-20 after each incubation. One hundred µl of TMB (3, 3', 5, 5' tetramethylbenzidine) substrate in peroxide solution was added to each well and incubated for 15 – 20 min. The reaction was stopped by adding 100 µl of 2M sulfuric acid and the solutions were read at 450 nm for absorbance, using a multiscan microplate reader (TiterTek, Huntsville, AL). The endpoint antibody titer is defined as the $\log_{10}$ value of the reciprocal of the highest serum dilution that yields an OD greater than the mean OD of sham-immunized control sera plus three standard deviations at the same serum dilution. The results are presented as the mean antibody endpoint titer and SE of the mean (SEM).

**Statistics**. The sample sizes for assaying vaccine efficacy post-challenge in hamsters (8/group for clinical symptomatology and oropharyngeal viral titers Days 1-3; 4/group for lung viral titer Day 3; 4/group for weight loss Days 1-7; 4/group for pathology Day 7) were estimated based on previous studies with 80% power using the usual alpha = 0.05 (i.e. $P < 0.05$) significance criterion. Body weight means on Days 3-7 and $\log_{10}$ PFU in the oropharyngeal swabs on Days 1-3 were compared among groups over time via repeated measure analysis of variance (ANOVA) models (mixed models). Mean $\log_{10}$ PFU in the cranial or caudal lungs on Day 3 and mean histopathological scores in cranial and caudal lungs on Day 7 were compared among groups using one way ANOVA. Normal quantile plots of the residual errors (not shown) confirm that the errors follow a normal distribution, as is required when using a parametric model. Analyses were carried out using R version 3.5.2 (R project for statistical computing) and JMP Pro version 15 (SAS Inc., Cary NC). A linear regression was used to obtain values for the slope and intercept and the correlation coefficient ($R^2$) between pre-challenge N protein specific serum IgG antibody endpoint titer and lung (cranial and caudal) histopathological scores at Day 7 post-challenge. Mean and standard error of the mean of serum antibody endpoint titer are reported, and means compared across groups by two-way analysis of variance (ANOVA) with Tukey's correction for multiple comparisons test using GraphPad Prism, 8.4.3 (San Diego, CA).

**Supplementary Materials and Methods**

**Generation of second generation multi-antigenic rLVS Δ*capB::*MN/SCoV2 vaccines.** We also generated second generation rLVS Δ*capB::*MN/SCoV2 vaccine candidates expressing the MN fusion protein from an antigen expression cassette integrated at the deleted *capB* locus in the chromosome and expressing the SΔTM, S1, or S2 protein from an antigen expression cassette located in a shuttle plasmid DNA. To construct rLVS Δ*capB* expressing the MN antigen from the chromosome, we amplified three DNA fragments - one encoding the antigen expression cassette Pbfr-N3F-MN, one homologous to the upstream region of the *capB* gene in the LVS Δ*capB* chromosome, and one homologous to the downstream region of the *capB* gene - by PCR using pFNL/Pbfr-N3F-MN and LVS Δ*capB* genomic DNA as templates for the first DNA fragment and for the second and third DNA fragments, respectively. We assembled the three DNA fragments by using the Gibson Assembly kit (NEB, Ipswich, MA), cloned it into the pMP590 integration plasmid [40] that contains a kanamycin resistance gene and a sucrose suicide gene, and verified the DNA sequence by restriction analysis and nucleotide sequencing. The resultant recombinant DNA, pMP/Pbfr-N3F-MN, was integrated into the *capB* locus by allelic exchange on Chocolate agar supplemented with kanamycin (7.5 µg/ml) followed by selection on Chocolate agar supplemented with 8% sucrose agar to generate marker-free rLVS Δ*capB*::MN. The chromosome integration of the MN expression cassette at the *capB* locus was verified by PCR for DNA integration and by Western blotting for MN fusion protein expression using monoclonal anti-FLAG M2 HRP antibody and/or guinea pig polyclonal antibody to SARS coronavirus. Subsequently, we introduced pFNL/Pbfr-N3F-SΔTM, pFNL/Pbfr-N3F-S1, and pFNL/Pbfr-S2 into rLVS Δ*capB*::MN to generate rLVS Δ*capB*::MN/SΔTM, S1 and S2 expressing MN plus SΔTM, MN plus S1, and MN plus S2, respectively. We verified these constructs for SARS-CoV-2 protein expression by Western blotting as described above.

**Immunogenicity study in mice.** Six to eight week old specific-pathogen-free female BALB/c mice were purchased from Charles River Laboratory (Sacramento, CA). The mice were immunized i.d. twice, 3 weeks apart, with normal saline (sham control), 2 x $10^6$ CFU LVS Δ*capB* (vector control), or 2 x $10^6$ CFU of rLVS Δ*capB*/SΔTM, S1, S2 or MN or with rLVS Δ*capB*::MN/SΔTM, S1 or S2. One week after the second immunization, mice were anesthetized by intraperitoneal injection of Ketamine (10 mg/ml) – Xylazine (1 mg/ml) solution, bled, and subsequently euthanized by inhalation of $CO_2$. Their spleens and lungs were removed and single cell suspensions of spleen and lung cells prepared as described by us previously [2,3]. Sera were isolated, heat inactivated, and frozen at -80$^{o}$C until use. T-cell mediated immune responses were examined by incubating single cell suspensions of lung and spleen cells with T-cell medium comprising Advanced RPMI-1640 (Invitrogen) supplemented with 2% heat-inactivated (HI) fetal bovine serum (Seradigm Premium Grade), penicillin (100 I.U./ml), streptomycin (100µg/ml), 0.1 mM non-essential amino acids, 4 mM L-glutamine, 1 mM sodium pyruvate, and 0.05 mM β-mercaptoethanol in the absence and presence of various SARS-CoV-2 and *F. tularensis* antigens; assaying for mouse interferon-gamma (IFN-γ); and quantitating intracellular cytokine staining by flow cytometry analysis. Humoral immune responses were examined by analyzing sera for levels of IgG and subtypes IgG1 and IgG2 antibodies specific for SARS-CoV-2 S RBD, N protein, and HI-LVS [41].

**Enzyme-linked immunosorbent assay (ELISA) for anti-SARS-CoV-2 antibody in mouse sera.** Mouse sera were assayed for IgG and subtype antibodies specific to SARS-CoV-2 S RBD and N protein antigens and to HI-LVS antigens similarly to what is described above for hamster serum with the following modifications. In addition to RBD and N protein antigens, wells were coated with SARS-CoV-2 S (1 μg/ml) or HI-LVS (5 x $10^6$/ml) diluted in carbonate/bicarbonate buffer. Sera at a starting dilution of 1:20 were diluted further through a three-fold series with PBS. After diluted sera were incubated with antigens coated on 96-well plates, the wells were incubated for 90 min with alkaline phosphatase (AP)-conjugated goat anti-mouse IgG (Sigma, St. Louis, MO), IgG1 or IgG2a (Invitrogen) at a dilution of 1:1000 at ambient temperature. After the plates were washed, 100 μl of NPP (*p*-nitrophenylphosphate) substrate in diethanolamine buffer (Phosphatase Substrate kit, BioRad, Hercules, CA) was added to each well and incubated for 15 – 20 min. The reaction was stopped by adding 100 μl of 0.1 N sodium hydroxide and the solutions were read at 415 nm for absorbance.

***In vitro* stimulation and production of IFN-γ by murine immune splenocytes**. A single cell suspension of 1.0 x $10^5$ splenocytes per well was seeded in U-bottom 96-well plates and incubated with T-cell medium alone, or T-cell medium supplemented with 2 μg/mL of recombinant SARS-CoV-2 protein or peptide antigens for three days. Afterwards, the culture supernatant fluid was collected, cell debris removed by centrifugation, and the supernatant fluid stored in assay diluent (BD Biosciences) at - 80°C until use. The production of mouse IFN-γ in the culture supernatant fluid was assayed using a mouse cytokine EIA kit (BD Biosciences).

***In vitro* stimulation and intracellular cytokine staining (ICS) for flow cytometry analysis.** A single cell suspension of 1.0 x $10^6$ splenocytes per well was seeded in U-bottom 96-well plates and stimulated with 2 μg/mL of recombinant S RBD or N protein or 5 x $10^6$/ml HI-LVS in the presence of anti-CD28 monoclonal antibody (Clone 37.51) for a total of 6 h, and processed for Flow Cytometry analysis as described previously by us [2]. Cells stimulated with medium alone (without antigen) or with PMA served as controls. The antibodies used for surface and intracellular cytokine staining included Alexa Fluor 488-conjugated hamster anti-mouse CD3e (clone 145-2c11), BV510-conjugated rat anti-mouse CD4 antibody (clone RM4-5), BV605-conjugated rat anti-mouse CD8 antibody (clone 53-6.7), BV711-conjugated rat anti-mouse IFN-γ (clone XMG1.2), BV421-conjugated anti-mouse TNF-α (clone MP6-XT22), PE-conjugated rat anti-mouse IL-2 (clone JES6-5H4), and Alexa Fluor 647-conjugated rat anti-mouse IL-17A antibody (clone TC11-18H10). All antibodies and reagents for ICS were purchased from BD Biosciences (San Jose, CA) unless otherwise indicated. The frequencies of live CD4+ and CD8+ T cells producing any of four cytokines (IFN-γ, TNF-α, IL-2, and IL-17A) were uniquely distinguished using FlowJo software. Background frequencies of cells producing cytokines without antigen stimulation were subtracted.

**Statistics**. The sample sizes for assaying immune responses post-vaccination in mice (4/group) were estimated based on previous studies with 80% power using the usual alpha = 0.05 (i.e. $p < 0.05$) significance criterion. Mean and standard error of the mean of serum antibody endpoint titer, cytokine production, and frequencies of cytokine-producing CD4+ and CD8+ T cells are reported, and means compared across groups by one-way or two-way analysis of variance (ANOVA) with Tukey's correction for multiple comparisons test using GraphPad Prism, 8.4.3 (San Diego, CA).

**Acknowledgement.** Flow cytometry was performed in the UCLA Jonsson Comprehensive Cancer Center (JCCC) and Center for AIDS Research Flow Cytometry Core Facility that is supported by National Institutes of Health awards P30 CA016042 and 5P30 AI028697, and by the JCCC, the UCLA AIDS Institute, and the David Geffen School of Medicine at UCLA.

### Supplemental Figures

Fig. S1

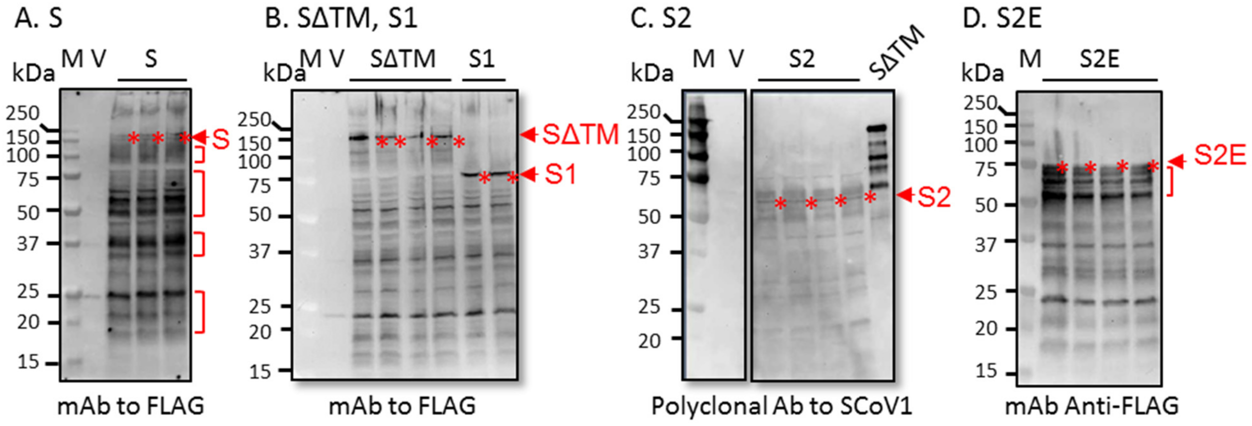


**Fig. S1. Expression of SARS-CoV-2 S, SΔTM, S1, S2, and S2E proteins by rLVS Δ*capB*/SCoV2 vaccines**. Expression of SARS-CoV-2 S (**A,** right 3 lanes), SΔTM (**B**, middle 4 lanes), S1 (**B**, right two lanes), S2 (**C**, right panel, left 4 lanes), and S2E (**D**, right 4 lanes) proteins by rLVS Δ*capB*/SCoV2 vaccines. Total bacterial lysates of LVS Δ*capB* vector (V, panels **A**, **B**, and **C**) and rLVS Δ*capB*/SCoV2 vaccine candidates (**A**, **B**, **C** right panel, and **D**) were analyzed by SDS-PAGE and Western blotting with an anti-FLAG monoclonal antibody (**A, B, D**) or an anti-SARS-CoV-1 guinea pig polyclonal antibody (BEI Resources, NR-10361) (**C**). The proteins of interest are indicated by red asterisks to the right of the protein band. The SARS-CoV-1 protein of SΔTM (BEI Resources, NR-722, ~150 kDa) (**C**, the rightmost lane) served as a positive control and was also detected by guinea pig polyclonal antibody against SARS-CoV-1. All panels: M, protein standards (lane 1); V, LVS Δ*capB* vector; the sizes of the molecular weight markers are labeled to the left of the panels.

Fig. S2

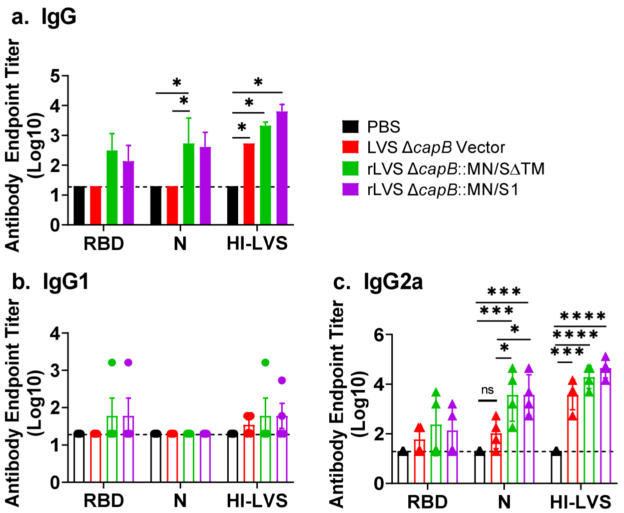


**Fig. S2. Humoral immune response induced by 2nd Generation rLVS Δ*capB*-vectored SARS-CoV2 vaccines in mice.** BALB/c mice (n=4/group) were sham-immunized or immunized ID twice at Week 0 and 3 with LVS Δ*capB* vector or 2nd generation rLVS Δ*capB*::MN/SCoV2/SΔTM and rLVS Δ*capB*::MN/SCoV2/S1 expressing the MN protein from the chromosome and the SΔTM and S1 proteins from a shuttle plasmid, as indicated to the right of the graph in **a**. At Week 4, mice were bled and sera assayed for IgG (**a**), IgG1 (**b**) and IgG2a (**c**) specific to SARS-CoV-2 S RBD or N protein or to heat-inactivated LVS Δ*capB* (HI-LVS). Data are mean log endpoint titer ± SE**.** *, P < 0.05, ***, P<0.001, ****P<0.0001 by Two-way ANOVA with Tukey's multiple comparisons (GraphPad Prism 8.02).

Fig. S3.

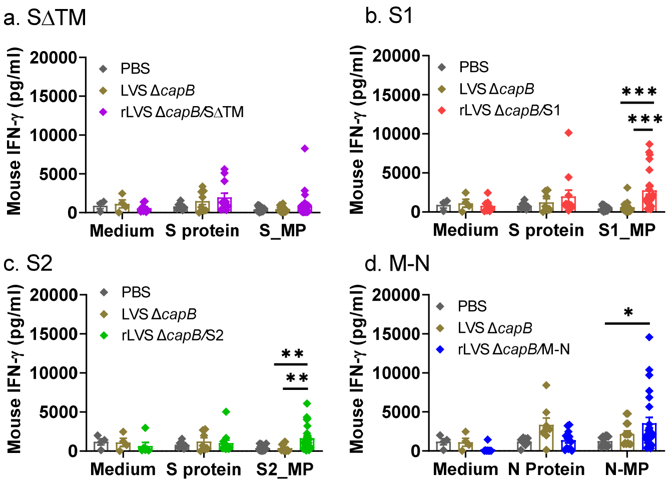


**Fig. S3. Cell-mediated immune responses induced by rLVS Δ*capB*-vectored SARS-CoV-2 vaccines in mice.** BALB/c mice (n=4/group) were immunized ID at Week 0 and 3; euthanized at Week 4; their splenocytes prepared, stimulated with proteins or master peptide pools (MP) in triplicate for 3 days; and culture supernatants collected and assayed for IFN-γ secretion. Data are mean IFN-γ secretion ± SE. *, $P < 0.05$; **, $P < 0.01$; ***, $P < 0.001$ by Two-way ANOVA with Tukey's multiple comparisons (GraphPad Prism 8.02).

**Supplemental Tables**

**Table S1. Key statistical analyses of impact of vaccines on weight loss and histopathology**

**A. Weight Loss**

| Comparison Groups | ID | IN | ID/IN |
|---|---|---|---|
| **Sham vs. MN** | <0.0001 | <0.01 | <0.0001 |
| **Sham vs. MN+SΔTM** | <0.004 | 0.001 | 0.0003 |
| **Sham vs. MN+S1** | 0.01 | 0.0002 | 0.0002 |
| **Sham vs. All MN Groups** | 0.0001 | 0.0001 | <0.0001 |
| **All MN vs. All S protein Groups** | 0.0008 | 0.0001 | <0.0001 |

**B. Histopathology - Combined Cranial and Caudal Lung Scores**

| Comparison Groups | ID | IN | ID/IN |
|---|---|---|---|
| **Sham vs. MN** | <0.0001 | <0.0001 | <0.0001 |
| **Sham vs. MN+SΔTM** | <0.0001 | <0.0001 | <0.0001 |
| **Sham vs. MN+S1** | <0.0001 | <0.0001 | <0.0001 |
| **Sham vs. All MN Groups** | <0.001 | <0.001 | <0.0001 |
| **All MN vs. All S protein Groups** | <0.001 | <0.001 | <0.0001 |

Means of each vaccine group Day 7 post-challenge were compared using a repeated measure (mixed) analysis of variance model. Normal quantile plots of the residual errors (not shown) confirm that the errors follow a normal distribution, as is required when using a parametric repeated measure model. ID, intradermal route of vaccination; IN, intranasal route of vaccination; ID/IN, intradermal or intranasal route of vaccination. All MN Groups, groups vaccinated with the rLVS Δ*capB*/SCoV2 vaccine expressing MN alone or combined with an rLVS Δ*capB*/SCoV2 vaccine expressing SΔTM or S1. All S Protein Groups, groups vaccinated with rLVS Δ*capB*/SCoV2 vaccine expressing S, SΔTM, S1, S2, or S2-E.

### Table S2. Lung Histopathology Scores

| | | Overall lesion | | | Pneumocyte | | Interstitial | |
|---|---|---|---|---|---|---|---|---|
| Hamster | Tissue[a)] | Extent | Bronchitis | Alveolitis | Hyperplasia | Vasculitis | Inflammation | Total score |
| Male | | | | | | | | |
| A3 | L1 | 3 | 2 | 4 | 4 | 2 | 4 | 19 |
| A4 | L1 | 4 | 2 | 5 | 5 | 2 | 5 | 23 |
| A7 | L1 | 3 | 1 | 4 | 4 | 2 | 4 | 18 |
| A8 | L1 | 3 | 1 | 4 | 4 | 2 | 4 | 18 |
| B3 | L1 | 3 | 1 | 4 | 3 | 1 | 4 | 16 |
| B4 | L1 | 3 | 1 | 4 | 3 | 3 | 4 | 18 |
| B7 | L1 | 3 | 2 | 4 | 4 | 3 | 4 | 20 |
| B8 | L1 | 3 | 2 | 4 | 4 | 2 | 4 | 19 |
| C3 | L1 | 4 | 2 | 5 | 4 | 2 | 5 | 22 |
| C4 | L1 | 3 | 1 | 4 | 4 | 2 | 4 | 18 |
| C7 | L1 | 3 | 2 | 4 | 4 | 2 | 4 | 19 |
| C8 | L1 | 3 | 2 | 4 | 4 | 2 | 4 | 19 |
| D3 | L1 | 4 | 2 | 5 | 4 | 2 | 5 | 22 |
| D4 | L1 | 3 | 0 | 3 | 2 | 2 | 3 | 13 |
| D7 | L1 | 3 | 2 | 4 | 4 | 2 | 4 | 19 |
| D8 | L1 | 3 | 1 | 4 | 4 | 1 | 4 | 17 |
| E3 | L1 | 3 | 2 | 4 | 4 | 2 | 4 | 19 |
| E4 | L1 | 4 | 3 | 5 | 4 | 2 | 5 | 23 |
| E7 | L1 | 3 | 2 | 4 | 4 | 1 | 4 | 18 |
| E8 | L1 | 3 | 1 | 4 | 4 | 1 | 4 | 17 |
| F7 | L1 | 3 | 2 | 4 | 4 | 1 | 4 | 18 |
| F8 | L1 | 3 | 2 | 4 | 4 | 1 | 4 | 18 |
| G3 | L1 | 3 | 2 | 4 | 4 | 2 | 4 | 19 |
| G4 | L1 | 3 | 2 | 4 | 3 | 3 | 4 | 19 |
| G7 | L1 | 3 | 2 | 4 | 4 | 3 | 4 | 20 |
| G8 | L1 | 3 | 2 | 4 | 4 | 2 | 4 | 19 |
| H3 | L1 | 1 | 0 | 1 | 0 | 0 | 2 | 4 |
| H4 | L1 | 1 | 0 | 0 | 1 | 0 | 2 | 4 |
| H7 | L1 | 1 | 0 | 1 | 1 | 0 | 1 | 4 |
| H8 | L1 | 1 | 0 | 1 | 0 | 2 | 2 | 6 |
| I3 | L1 | 1 | 2 | 0 | 0 | 2 | 2 | 7 |
| I4 | L1 | 1 | 0 | 0 * | 0 | 0 | 2 | 3 |
| I5 | L1 | 1 | 0 | 1 | 0 | 3 | 2 | 7 |
| I7 | L1 | 2 | 1 | 1 | 0 | 4 | 2 | 10 |
| J4 | L1 | 2 | 0 | 2 | 2 | 2 | 2 | 10 |
| J7 | L1 | 2 | 2 | 0 | 0 | 3 | 2 | 9 |
| J8 | L1 | 2 | 2 | 2 | 0 | 3 | 2 | 11 |
| K3 | L1 | 3 | 1 | 4 | 4 | 2 | 4 | 18 |
| K4 | L1 | 3 | 1 | 4 | 4 | 1 | 4 | 17 |
| K7 | L1 | 3 | 2 | 4 | 4 | 2 | 4 | 19 |
| K8 | L1 | 3 | 1 | 4 | 4 | 1 | 4 | 17 |
| L3 | L1 | 3 | 1 | 4 | 4 | 2 | 4 | 18 |
| L4 | L1 | 3 | 2 | 4 | 4 | 1 | 4 | 18 |
| L7 | L1 | 3 | 2 | 4 | 4 | 2 | 4 | 19 |
| L8 | L1 | 3 | 2 | 4 | 4 | 2 | 4 | 19 |

**Table S2 Continued**

| | | Overall lesion | | | Pneumocyte | | Interstitial | |
|---|---|---|---|---|---|---|---|---|
| Hamster | Tissue[a)] | Extent | Bronchitis | Alveolitis | Hyperplasia | Vasculitis | Inflammation | Total score |
| M3 | L1 | 3 | 1 | 3 | 3 | 2 | 3 | 15 |
| M4 | L1 | 3 | 1 | 4 | 4 | 2 | 4 | 18 |
| N3 | L1 | 4 | 2 | 5 | 5 | 1 | 5 | 22 |
| N4 | L1 | 3 | 2 | 4 | 4 | 2 | 4 | 19 |
| N7 | L1 | 3 | 2 | 4 | 4 | 2 | 4 | 19 |
| N8 | L1 | 3 | 2 | 4 | 4 | 3 | 4 | 20 |
| O3 | L1 | 4 | 1 | 5 | 5 | 2 | 5 | 22 |
| O4 | L1 | 2 | 0 | 3 | 3 | 0 | 3 | 11 |
| O8 | L1 | 3 | 1 | 4 | 4 | 2 | 4 | 18 |
| P3 | L1 | 3 | 2 | 4 | 4 | 1 | 4 | 18 |
| P4 | L1 | 3 | 1 | 4 | 4 | 1 | 4 | 17 |
| P7 | L1 | 3 | 2 | 4 | 4 | 3 | 4 | 20 |
| P8 | L1 | 3 | 2 | 4 | 4 | 1 | 4 | 18 |
| Q3 | L1 | 2 | 0 | 0 | 0 | 0 | 2 | 4 |
| Q4 | L1 | 2 | 0 | 1 | 0 | 3 | 2 | 8 |
| Q7 | L1 | 1 | 0 | 1 | 1 | 0 | 1 | 4 |
| Q8 | L1 | 2 | 2 | 2 | 0 | 2 | 2 | 10 |
| R3 | L1 | 2 | 1 | 2 | 2 | 1 | 2 | 10 |
| R4 | L1 | 2 | 0 | 1 | 1 | 2 | 2 | 8 |
| R7 | L1 | 2 | 2 | 2 | 2 | 2 | 2 | 12 |
| R8 | L1 | 2 | 1 | 2 | 2 | 2 | 2 | 11 |
| S4 | L1 | 1 | 0 | 0 | 0 | 1 | 1 | 3 |
| S7 | L1 | 1 | 1 | 0 | 0 | 3 | 2 | 7 |
| S8 | L1 | 2 | 1 | 3 | 3 | 2 | 3 | 14 |
| Female | | | | | | | | |
| A3 | L2 | 4 | 2 | 4 | 4 | 3 | 4 | 21 |
| A4 | L2 | 3 | 2 | 4 | 4 | 2 | 4 | 19 |
| A7 | L2 | 3 | 1 | 4 | 4 | 1 | 4 | 17 |
| A8 | L2 | 3 | 2 | 4 | 4 | 2 | 4 | 19 |
| B3 | L2 | 3 | 2 | 4 | 4 | 2 | 4 | 19 |
| B4 | L2 | 4 | 1 | 5 | 5 | 1 | 5 | 21 |
| B7 | L2 | 3 | 2 | 4 | 4 | 3 | 4 | 20 |
| B8 | L2 | 3 | 2 | 4 | 4 | 2 | 4 | 19 |
| C3 | L2 | 3 | 2 | 4 | 3 | 2 | 4 | 18 |
| C4 | L2 | 3 | 2 | 4 | 4 | 2 | 4 | 19 |
| C7 | L2 | 3 | 2 | 4 | 4 | 3 | 4 | 20 |
| C8 | L2 | 3 | 2 | 4 | 4 | 2 | 4 | 19 |
| D3 | L2 | 4 | 2 | 4 | 4 | 2 | 4 | 20 |
| D4 | L2 | 3 | 2 | 4 | 4 | 2 | 4 | 19 |
| D7 | L2 | 3 | 2 | 4 | 4 | 2 | 4 | 19 |
| D8 | L2 | 3 | 3 | 4 | 4 | 2 | 4 | 20 |
| E3 | L2 | 3 | 1 | 4 | 4 | 2 | 4 | 18 |
| E4 | L2 | 4 | 3 | 5 | 4 | 2 | 5 | 23 |
| E7 | L2 | 3 | 2 | 4 | 4 | 2 | 4 | 19 |
| E8 | L2 | 3 | 0 | 4 | 4 | 2 | 4 | 17 |
| F7 | L2 | 3 | 1 | 4 | 4 | 2 | 4 | 18 |
| F8 | L2 | 3 | 2 | 4 | 4 | 2 | 4 | 19 |

**Table S2 Continued**

| | | Overall lesion | | | Pneumocyte | | Interstitial | |
|---|---|---|---|---|---|---|---|---|
| **Hamster** | **Tissue[a)]** | **Extent** | **Bronchitis** | **Alveolitis** | **Hyperplasia** | **Vasculitis** | **Inflammation** | **Total score** |
| G3 | L2 | 4 | 3 | 5 | 4 | 3 | 5 | 24 |
| G4 | L2 | 4 | 2 | 5 | 4 | 4 | 5 | 24 |
| G7 | L2 | 2 | 2 | 4 | 4 | 2 | 4 | 18 |
| G8 | L2 | 3 | 2 | 4 | 4 | 4 | 4 | 21 |
| H3 | L2 | 1 | 0 | 0 | 0 | 0 | 2 | 3 |
| H4 | L2 | 1 | 0 | 2 | 2 | 1 | 2 | 8 |
| H7 | L2 | 1 | 0 | 2 | 2 | 0 | 2 | 7 |
| H8 | L2 | 2 | 2 | 1 | 0 | 1 | 2 | 8 |
| I3 | L2 | 1 | 0 | 0 | 0 | 3 | 2 | 6 |
| I4 | L2 | 2 | 0 | 1 | 0 | 0 | 2 | 5 |
| I5 | L2 | 1 | 0 | 0 | 0 | 3 | 2 | 6 |
| I7 | L2 | 1 | 0 | 0 | 0 | 3 | 2 | 6 |
| J4 | L2 | 1 | 1 | 1 | 2 | 1 | 2 | 8 |
| J7 | L2 | 2 | 0 | 0 | 0 | 2 | 2 | 6 |
| J8 | L2 | 2 | 1 | 2 | 1 | 3 | 2 | 11 |
| K3 | L2 | 3 | 1 | 4 | 3 | 1 | 4 | 16 |
| K4 | L2 | 3 | 1 | 4 | 4 | 1 | 4 | 17 |
| K7 | L2 | 3 | 2 | 3 | 3 | 1 | 3 | 15 |
| K8 | L2 | 3 | 1 | 4 | 4 | 2 | 4 | 18 |
| L3 | L2 | 3 | 1 | 4 | 4 | 2 | 4 | 18 |
| L4 | L2 | 2 | 1 | 3 | 3 | 2 | 3 | 14 |
| L7 | L2 | 3 | 1 | 4 | 4 | 2 | 4 | 18 |
| L8 | L2 | 3 | 2 | 4 | 4 | 1 | 4 | 18 |
| M3 | L2 | 3 | 1 | 4 | 4 | 1 | 4 | 17 |
| M4 | L2 | 3 | 1 | 4 | 4 | 1 | 4 | 17 |
| N3 | L2 | 3 | 1 | 4 | 4 | 2 | 4 | 18 |
| N4 | L2 | 4 | 2 | 5 | 4 | 3 | 5 | 23 |
| N7 | L2 | 3 | 1 | 4 | 4 | 1 | 4 | 17 |
| N8 | L2 | 3 | 2 | 4 | 4 | 1 | 4 | 18 |
| O3 | L2 | 3 | 2 | 4 | 4 | 3 | 4 | 20 |
| O4 | L2 | 3 | 2 | 4 | 4 | 2 | 4 | 19 |
| O8 | L2 | 3 | 1 | 4 | 4 | 2 | 4 | 18 |
| P3 | L2 | 4 | 1 | 4 | 4 | 2 | 4 | 19 |
| P4 | L2 | 3 | 2 | 4 | 4 | 0 | 4 | 17 |
| P7 | L2 | 3 | 2 | 4 | 4 | 2 | 4 | 19 |
| P8 | L2 | 3 | 1 | 4 | 4 | 2 | 4 | 18 |
| Q3 | L2 | 2 | 0 | 2 | 2 | 0 | 2 | 8 |
| Q4 | L2 | 1 | 0 | 1 | 2 | 3 | 2 | 9 |
| Q7 | L2 | 1 | 0 | 2 | 1 | 0 | 1 | 5 |
| Q8 | L2 | 2 | 1 | 2 | 2 | 0 | 2 | 9 |
| R3 | L2 | 3 | 2 | 3 | 3 | 1 | 3 | 15 |
| R4 | L2 | 2 | 0 | 2 | 2 | 2 | 2 | 10 |
| R7 | L2 | 2 | 2 | 3 | 3 | 1 | 3 | 14 |
| R8 | L2 | 1 | 0 | 1 | 1 | 0 | 1 | 4 |
| S4 | L2 | 2 | 1 | 0 | 0 | 3 | 3 | 9 |
| S7 | L2 | 1 | 1 | 0 | 0 | 2 | 1 | 5 |
| S8 | L2 | 2 | 0 | 3 | 2 | 1 | 3 | 11 |

a) L1, cranial; L2, caudal
